# The microtubule organization in the zebrafish yolk adapts to transgene-mediated phenotypic variations

**DOI:** 10.1101/2020.08.06.239970

**Authors:** Maria Marsal, Matteo Bernardello, Emilio J. Gualda, Pablo Loza-Alvarez

## Abstract

The organization of microtubule networks in the cells is orchestrated by subcellular structures named microtubule organizing centers (MTOCs). In zebrafish embryos, the yolk is surrounded by a cytoplasmic layer containing a vast network of microtubules. In order to understand how this complex network is organized, we use dclk2-GFP zebrafish embryos, as a microtubule reporter line, and Light Sheet Fluorescence Microscopy. We find that this organization is mediated by a variable number of aster-like MTOCs during epiboly, and that it does not follow a rigid scheme, exemplifying developmental robustness. We characterize asters morphology, dynamics, and their uniform distribution in the yolk sphere. Consistent with their role as MTOCs we find that they contain key molecular machinery for MTs dynamics, amongst which centrin marks the assignation of MTOCs over time. Finally, we demonstrate that merely the overexpression of dclk2-GFP in wild type embryos can induce the formation of asters. We propose dclk2-GFP embryos as a model for the study of the collective behaviour of microtubules in complex systems.

## INTRODUCTION

Epiboly is one of the cell movements of gastrulation, highly conserved throughout evolution. It consists of the thinning and spreading of different cell layers over and around the yolk cell (Betchaku and Trinkaus, 1978; Warga and Kimmel, 1990). In zebrafish, after fertilization, meroblastic cleavages give rise to the blastoderm, sitting on top of the big yolk cell that does not divide. After the 9^th^ or 10^th^ division cycle, the more marginal blastoderm cells release their cell contents into the yolk, forming a syncytium (Chu *et al*., 2012). This is named the yolk syncytial layer (YSL), which is adjacent to a thin cytoplasmic layer (YCL) that wraps the viscous yolk mainly composed of lipid granules (Kimmel and Law, 1985; Kimmel *et al*., 1995; Carvalho and Heisenberg, 2010). The nuclei of the YSL (YSN) undergo three rounds of divisions and become post-mitotic (Kane, Warga and Kimmel, 1992). Soon after that, some YSN move underneath the blastoderm (internal-YSN, i-YSN) and others stay at the surface (external-YSN, e-YSN) and undergo epiboly together with the blastoderm (D’Amico and Cooper, 2001; Carvalho *et al*., 2009). Acto-myosin motors in the YSL and a polarized gradient of cortical tension are currently seen as the driving epiboly motors (Cheng, Miller and Webb, 2004; Köppen *et al*., 2006; Behrndt *et al*., 2012; Bruce, 2016; Hernández-Vega *et al*., 2017).

It is known, that the zebrafish YSL and YCL contain microtubule (MT) networks that undergo various changes (MT density, MT length) over the different developmental stages (Solnica-Krezel and Driever, 1994; Jesuthasan and Strähle, 1997; Tran *et al*., 2012; Eckerle *et al*., 2018). However, during embryogenesis, little is known about the specific mechanisms that pattern such a network. In general, the organization of MTs is not random but is orchestrated by defined subcellular regions called MT organizing centers (MTOCs). Often, the MTs are organized in a radial way, nucleated from a central organizing region, forming what it is known as a MT aster. This is how the centrosomes, used by dividing animal cells, organize the mitotic spindle. The best-studied MTOC is the centrosome but it has further been shown that differentiated cells can generate alternative MT organizations through the reassignment of MTOC functions to non-centrosomal sites in interphase cells. These non-centrosomal MTOCs (nc-MTOCs) often serve as mechanical support or intracellular transport scaffolds (Sanchez and Feldman, 2017).

In zebrafish, yolk cell MTs are necessary for the migration of the e-YSN (Solnica-Krezel and Driever, 1994; Fei *et al*., 2019). However, their role, either instructive or permissive is not yet fully understood. Yolk cell MTs are sensitive to environmental conditions (Strahle and Jesuthasan, 1993; Jesuthasan and Strähle, 1997; Sarmah *et al*., 2013) and the defects in the yolk MT network are often seen in epiboly mutants. However, it is difficult to know if the MT defects are the primary cause of the phenotype (Du et al., 2012; (Li-Villarreal *et al*., 2015; Eckerle *et al*., 2018). Not only the function but also the arrangement of the MT network in the zebrafish yolk cell is not yet completely understood. This is because the problem has been approached only partially, both in time and in space. In fact, the primary approach has been immunostaining on fixed samples (Strahle and Jesuthasan, 1993; Topczewski and Solnica-Krezel, 1999), thus, hiding the rich variety of phenotypes over time and its dynamic nature. Only recently, the use of reporter transgenic lines and laser scanning confocal microscopy allowed the dynamical study of yolk cell MTs, although only in restricted areas and in a limited time window (Tran *et al*., 2012; Eckerle *et al*., 2018; Fei *et al*., 2019).

In this work, we use Light-Sheet Fluorescence Microscopy (LSFM), Laser Scanning Confocal Microscopy (LSCM) and the MT reporter line Tg(Xla.Eef1a1:dclk2a-GFP) (Tran *et al*., 2012) to obtain a mesoscopic description of the spatio-temporal yolk cell MT organization during early zebrafish development and to find out the cellular and molecular mechanism responsible of building this network.

Thanks to the high-throughput capabilities of our custom-made LSFM microscope we are now able to quantitatively assess the high phenotypic variability of yolk MT organization. Thus, in addition to confirming the existence of previously reported MTOCs in the YSL (Solnica-Krezel and Driever, 1994; Fei *et al*., 2019) we provide new insights on the mesoscopic description of the yolk MT organization and identify, at middle-stage epiboly and for the first time, the presence of MTOCs in the YCL. Here, we perform a thorough analysis of this newly find yolk MT network organization. We find that the YCL MTOCs appear regularly in a precise time window occupying the yolk surface, and assemble into large 3D patterns of radially oriented MT (YCL asters). We analyze their spatial dimensions and microns-scale architecture and hypothesize on how YCL asters interact at common boundaries. We quantify their position and variable number in the transgenic embryos, where YCL asters group in one or more spherical latitudes. This variability thus encodes different configurations, compatible with development, highlighting the yolk domain as a plastic but robust region. We also manage to induce the formation of YCL asters in wild type embryos upon the transient expression of the MT associated proteins DCLK (doublecourtin-like kinase) or DCX (doublecourtin), important for MT stabilization and nucleation (Moores *et al*., 2006; Ramkumar, Jong and Ori-McKenney, 2018). We address how the presence and number of YCL asters impact on epiboly progression.

As LSFM is well suited to assess the yolk MT organization in the embryo as a whole, we manage to link different MTOCS in time and in space and to study the overall distribution of MTOCs in the yolk sphere. The large syncytial yolk cell (~700-800μm) is similar in size to amphibian zygotes (~1200μm) and far exceeding the size of standard somatic cells. Therefore, it is an ideal model for studying MT based spatial organization in big cells. We propose that the mechanism (i.e the formation of multiple MT asters) shown here could exemplify a special adaptation of conserved mechanisms. In addition, it allows to underscore unnoticed capabilities of the yolk cell.

Finally, we also identify the key molecules of MTOCs in the zebrafish yolk cell. Amongst them, we determine that centrin highlights both centrosomal and non-centrosomal MTOCs in the yolk cell and allows us to follow the potential reassignment of new sites as MTOCs over time.

## RESULTS

### The yolk MT network organizes into different microtubule organizing centers (MTOCs) during early zebrafish development

Thanks to our custom-made high throughput multi-view light sheet imaging approach, we have access, with high temporal and spatial resolution, to the whole embryonic sphere throughout zebrafish early developmental stages. LSFM provides high resolution with high signal to noise ratio images in a very efficient way. This allows visualizing even the internal structures of the yolk sphere, with less photo-bleaching and photo-toxicity (Olarte et al., 2018; Andilla et al., 2017). Additionally, our original high-throughput sample mounting solution based on fluidic sample loading (Gualda et al., 2015) permitted the study of a large population (over one hundred) of embryos preserving sample viability and in a highly reproducible way. This has allowed us to show the great phenotypic variability on the yolk MT network. Some examples are displayed in Figure 1.

**Figure 1.**
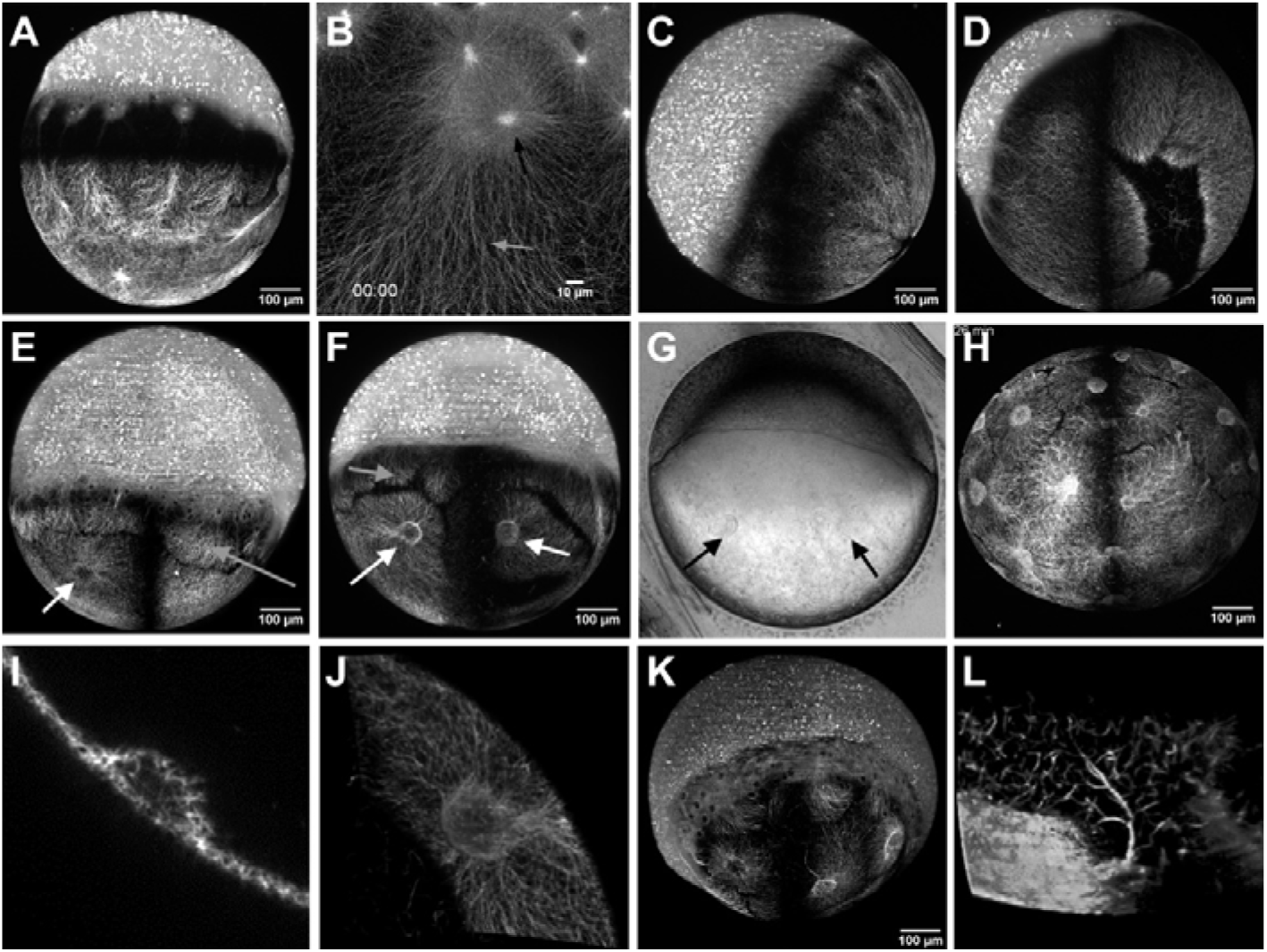
Variability on yolk microtubule organization centers (MTOCs) in dclk2-GFP transgenic zebrafish embryos using light sheet fluorescence microscopy. **(A)** At sphere stage, parallel MT arrays emerge from YSL, covering the YCL. At the animal pole, mitotic spindles of dividing cells are also visible. **(B)** LSCM zoom into the YSL, we can see meshed interconnected MT with apparent spindles during mitosis of e-YSN (black arrow) and the AV parallel YCL MT arrays, emanating from MTOCs associated to the most vegetal e-YSN (grey arrow). **(C)** At 50 % epiboly, the blastoderm occupies half of the embryo sphere. **(D)** Embryo showing defined MT domains and aperture of the vegetal pole. **(E)** Many embryos present YCL asters, a radial MT organization in defined regions (white arrows). A cross section of each YCL aster **(I)** shows an invagination of the yolk membrane, covered with MT. **(F)** These YCL asters organize MT in clear domains, different from the MT network emerging from the YSL (grey arrows). A volume render **(J)** helps highlighting the 3D nature of these structures, also visible in **(G)** bright-field imaging (black arrows). **(H)** Some embryos show up to 22 YCL asters, covering the entire YCL. In many asters, thicker MT bundles are visible in specific directions. **(K)** Most of the embryos showing YCL asters lack a vegetal pole aperture. In some cases, an aster occupies this position, and occasionally there is a connection to thick MT bundles from the inner MT network **(L)**. All the described features can be clearly observed in the 3D volume rendering in Video S1.

We have primarily used Tg(Xla.Eef1a1:dclk2a-GFP) (from now on abbreviated dclk2-GFP) embryos, as a MT reporter transgenic line. Through the dclk2-GFP line we can observe, before epiboly starts, the presence of the two types of yolk MTs organization already described in the 90’s (Solnica-Krezel and Driever, 1994; Jesuthasan and Strähle, 1997). Those are, firstly, meshed interconnected MTs covering the entire YSL corona, with apparent spindles during mitosis of YSN (Figure 1B). Secondly, the animal-vegetal (AV) parallel arrays of MTs of the YCL, emanating from potential MTOCs associated to the most vegetal e-YSN and ending at different latitudes of the yolk cell (Figures 1A and 1B) (Jesuthasan and Strähle, 1997; Eckerle *et al*., 2018).

At 50% epiboly, when the deep cells engage in the gastrulation movements of involution and convergence, the blastoderm covers almost completely the YSN and their associated MT arrays (Figure 1C). From this point, the possibility to inspect many embryos with our LSFM microscope revealed, despite the classical view, a large phenotypic variability in the MT organization along the yolk cell. While some embryos show clearly the AV parallel arrays, in others these arrays are grouped in fewer well-defined domains. Moreover, some embryos show an opening on the vegetal pole (Figure 1D), while others do not. Finally, and common to most of the analyzed embryos, the vegetal array of MTs organizes in clear and visible compartments that we will call yolk cytoplasmic layer asters (YCL asters) (Figures 1E, 1F, 1H and 1K), with MTs radially orienting from each of the domains center. To the best of our knowledge, this is the first time that those structures are reported. Interestingly, a simple observation with bright-field microscopy also reveals invaginations of the yolk membrane at the sites of each YCL aster (Figure 1G). The number of the YCL asters can vary from 0 up to 22 in a single embryo (Figure 1H). A cross-section (Figure 1I) reveals that those YCL asters present a double MT network, one at the yolk surface and another surrounding basally a region with less MT density. A 3D render (Figure 1J and Video S1) shows how those structures resemble a half sphere shape. Remarkably, in some cases, thicker MT bundles from deep inside the yolk mass assemble and extend from the bottom of the asters in a process of MT recruitment (Figures 1K, 1L and S1).

Interestingly, in eggs with many YCL asters, visual inspection suggests that their distribution is not random but occurs in one or more concentric rings around the animal-vegetal axis. To confirm this observation, we calculated the spherical coordinates of the YCL asters. Results are displayed in Figure 2, where we show two representative embryos with four rings (Figure 2 A-D) and a single ring (Figure 2 E-H), respectively. The computation of their polar angles (latitude coordinates, ⍰) confirms that different asters indeed group in the same latitude (Figures 2B, 2C, 2F and 2G). In embryos with more than one concentric ring, different latitudes can be observed (Figure 2B and 2C). The number of rings in the different embryos is variable, as for the number of YCL asters in a ring (Figures 2I and 2J). Finally, the computation of the aster’s azimuthal angles (longitude coordinates, φ) (Figures 2D and 2H) showed that asters within a ring are equidistant in a range between 37.5% to 100%, independently of the total number of asters in the ring (Figure 2K). The yolk view was not always complete due to the embryo position, therefore not all YCL asters were visible. This fact could account for the non-equidistancy portion. In those cases (see asteriks in figure 2D) the distances are doubled since most probably an aster in between was not imaged.

**Figure 2.**
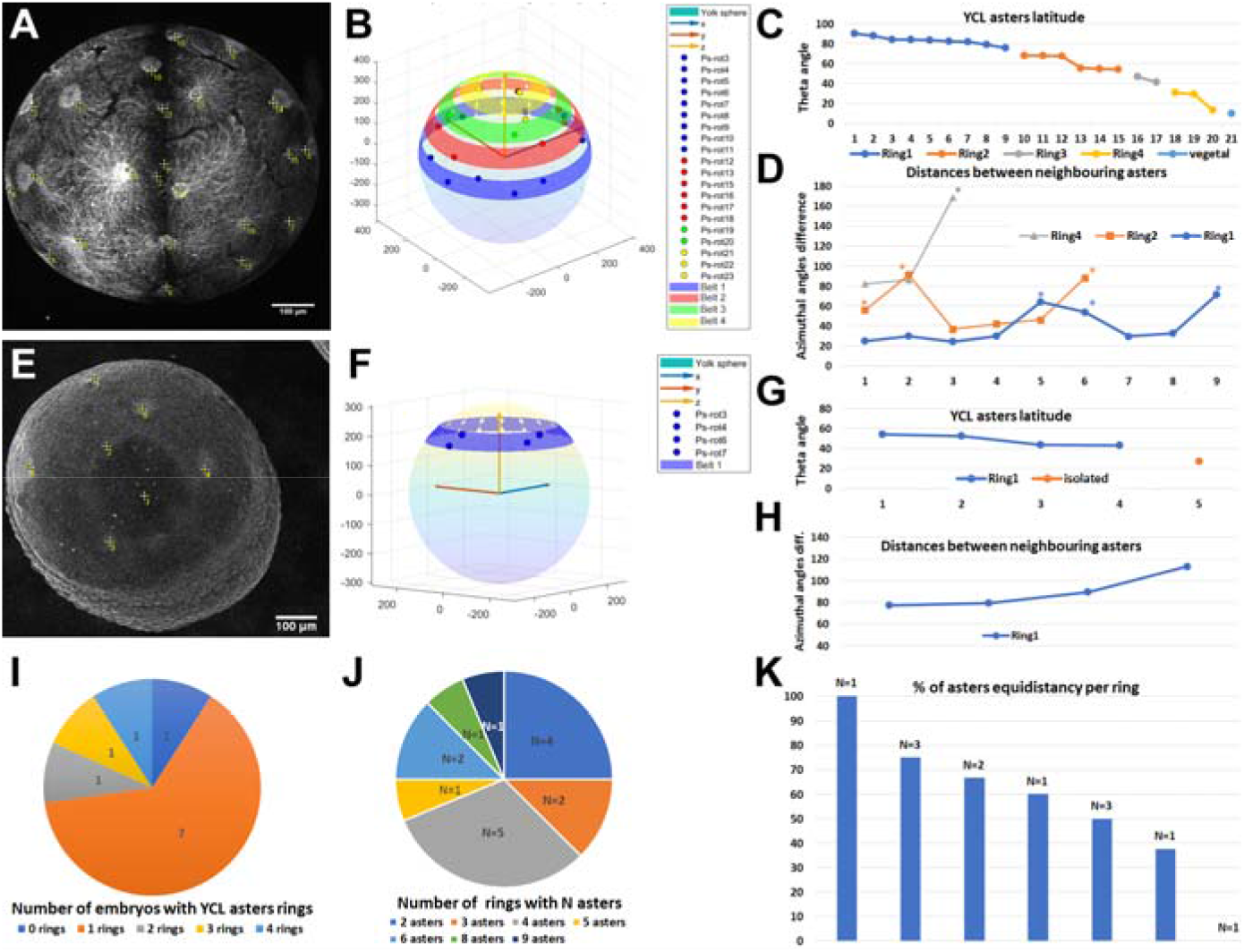
Analysis of the spatial YCL aster distribution in rings. **(A)** Maximum projection of a dclk2-GFP embryo where 22 YCL asters can be identified (yellow crosses) distributed in four rings. **(B)** Projection of the aster position in a sphere. Asters belonging to the same ring are represented by dots of the same color, rings are highlighted as belts at the calculated latitude. **(C)** YCL asters latitude (polar angle). YCL asters are not randomly located but group in four different latitudes (rings) (blue, orange, grey and yellow) and a single aster is found at vegetal pole. **(D)** Azimuthal angles difference between neighboring asters in rings 1 (blue), 2 (orange) and 4 (grey). Asters in this embryo are equidistant in 11/18 cases. For cases they are not (asterisk), we suspect an aster in between could not be imaged. **(E-H)** Same analysis for an embryo with only one YCL asters ring. Here 5 YCL asters can be identified. For this particular case, asters are equidistant: 80, 80, 90 and 110 (5 asters distributed around 360 degrees). **(I)** Plot of the distribution of YCL asters in 0, 1, 2, 3 or 4 rings. The analysis was performed in 11 embryos. **(J)** Quantification of the number of asters in different rings. The analysis was performed in 16 rings. **(K)** Quantification of the equidistance between YCL asters, analyzed in 12 rings. Regardless the total YCL aster number per ring, equidistance ranged between 37.5% to 100%. Only in one case no equidistance was observed. N, number of rings.

### YCL asters are transient 3D patterns of radially oriented MTs

To gain insight on the dynamics and the architecture of YCL asters, we next examined them with high temporal and spatial resolution. We could split the temporal YCL asters evolution in four steps: compartmentalization of yolk MTs into distinguishable domains; formation of asters with MTs radially orienting from the center of each compartment; growth of asters; and finally, asters reabsorption below the approaching blastoderm.

The re-organization of the yolk MT network that gives rise to this scenario begins when the YSN become post-mitotic, that is, when epiboly starts. We clearly observe a transition between a uniformly yolk coverage of MTs (Figure 3A) to the formation of domains (Figure 3B), that look independent from each other at this scale of observation. At this point, emerging from the blastoderm, YCL asters form as they migrate vegetally along the MT bundles anchored at the YSL (Figure 3C). At 65% epiboly, just after the formation of the embryonic shield (Warga and Kimmel, 1990; Montero *et al*., 2005), the e-YSN re-emerge from below the blastoderm and start moving towards the vegetal pole. At this moment, all the previously formed vegetal MTs compartments generate a clearly distinguishable round central region with MT bundles that are radially oriented from it (Figure 3D). YCL asters span the yolk surface and resemble large MTOCs. The degree of epiboly when they form can vary between embryos and, in some cases, the e-YSN mitotic asters coexist with these newly formed YCL asters (Video S2). In rare cases, we observe the formation of a dynamic vortex-like organization of the yolk MTs prior to the formation of the YCL asters (Video S3), very similarly to *in vitro* reconstitution models (Juniper *et al*., 2018).

**Figure 3.**
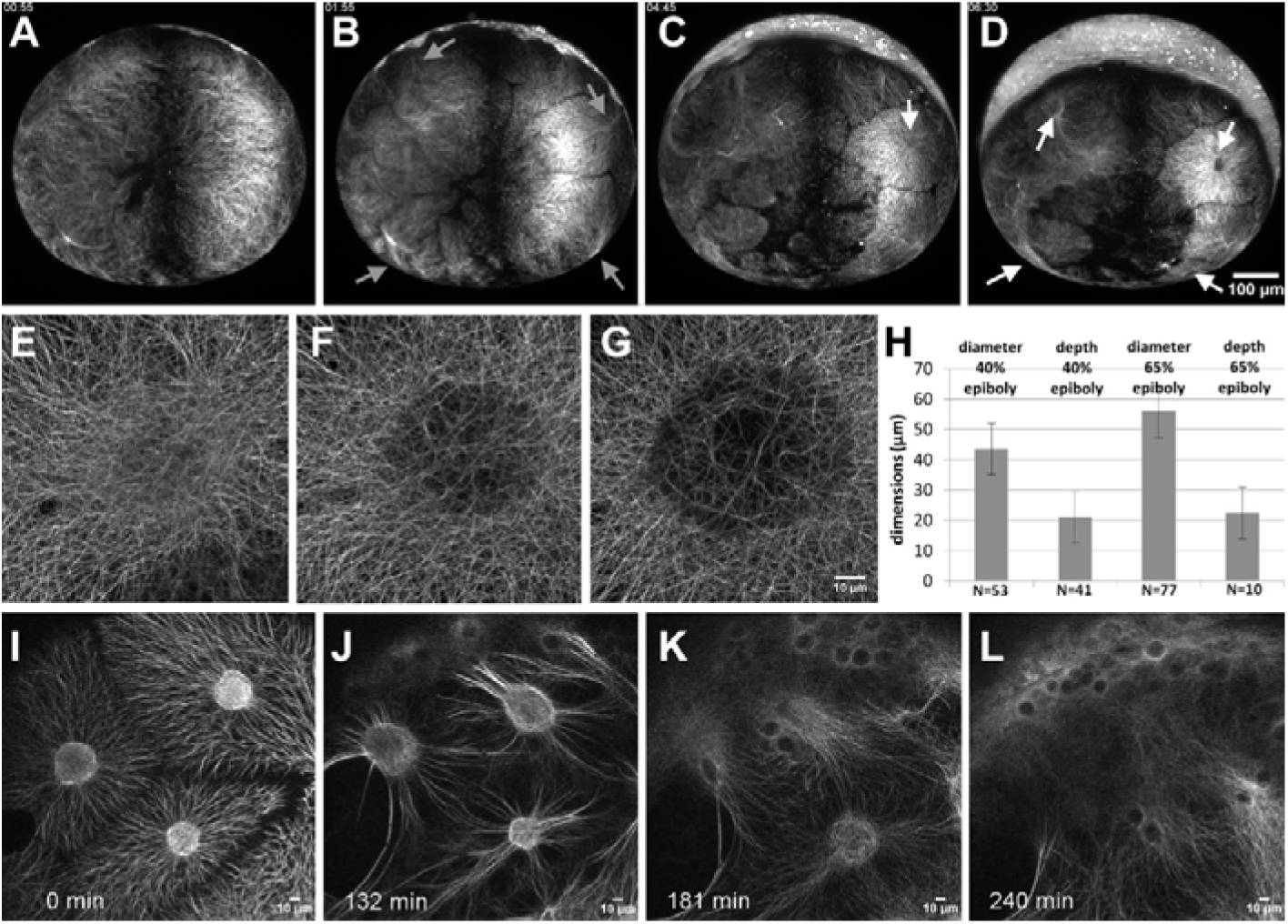
The YCL asters form and evolve throughout epiboly. **(A)** After the last YSN division, the MT network acquires a new configuration. **(B)** At sphere stage, the MT network has rearranged into clear domains, with a central high dense MT bundle emanating from the YSL MTOCs (grey arrow). **(C)** Emerging from those bundles, defined MTs domains start migrating vegetally following a flat to hemisphere transition, leading to the formation of a visible YCL aster (white arrow) with radially oriented MT fibers in the middle of each domain **(D)**. **(E-G)** High-resolution visualization of an aster growth through LSCM. (H) After inspection of more than hundred asters in different embryos we observe that, once formed, the depth of the asters keeps constant while their diameter increases. **(I)** The aster domains remain individualized, with clear opposing MT tips. In some cases, the YCL aster core shows a brighter fluorescence signal, while others appear emptier **(G).** When the marginal blastoderm approaches the YCL asters, the MT network rearranges once again. Thicker bundles are formed and reorient in the AV direction (J) and the cores of the YCL asters actively migrate animalwards (K) until they disappear underneath the YSL **(L).** See also Videos S2 and S6.

In Figures 3E-G and Video S4 we show, using high resolution LSCM, the initial steps of YCL aster formation. Before an actual YCL aster is formed, a dense central circle of disorganized MTs can be identified (Figure 3E). Later on, the density of MTs is slowly reduced in this central region evidenced by showing a reduced signal (Figure 3F). Progressively a membrane depression forms the final YCL aster (Figure 3G). At this point the surrounding dense network of MTs orients radially from the center of the aster and clear MT domains can be distinguished.

A 3D analysis shows that they change over time from a flat shape to a half sphere shape that is fully filled with thicker MTs bundles. We performed a statistical analysis of the diameter, depth and distribution of more than 80 YCL asters from different embryos. On average, the hemispheres display a maximum diameter of 43 μm at 40% epiboly and 56 μm at 65% epiboly, with a constant average depth of 21-22μm from the base to the vertex (Figure 3H). Zooming into the interaction area between asters, we observe clear boundaries between YCL asters domains (Figure 3I) where MTs seem to repel each other.

Once formed, the YCL asters persist on a quasi-static state and in the same position at the mesoscopic level of observation. Only when the blastoderm undergoing epiboly approaches, YCL asters dynamically move beneath the blastoderm changing their shape (Figures 3I-L and Video S5). We can observe a general displacement of the YCL aster towards the animal pole and a reorganization of the MTs in the aster, from a more uniform distribution to a biased preferred distribution (parallel to the AV direction of epiboly movement) (Figures 3I, 3J and S2). Soon after, the asters interact with the migrating e-YSN at the same time they are reabsorbed beneath the blastoderm (Figures 3K, 3L and Video S6).

As described above, in embryos with many asters we have noticed that the asters organize uniformly forming rings on the YCL at specific latitudes. In those, we observed the synchronous reabsorption of the asters belonging to the same ring. Rings of asters laying towards the more vegetal pole are sequentially reabsorbed, triggered by the proximity of the YSL and blastoderm undergoing epiboly.

### MT polymerization and nucleation occur at YSN centrosomes and YCL asters

The above observations suggest that YCL asters could be acting as MTOCs. MTOCs are specific sites where MTs are nucleated and from where filaments grow through polymerization of α- and β-tubulin dimers (Lüders and Stearns, 2007; Wu and Akhmanova, 2017). Probably, the most well-known MTOC is the centrosome used by cells during mitosis. However, in many non-dividing cells, the organization of the MTs is imparted by non-centrosomal MTOCs (nc-MTOCs), whose composition is rather unknown (Muroyama and Lechler, 2017; Sanchez and Feldman, 2017). Nc-MTOCs are thought to contain a shorter list of proteins compared to centrosomal MTOCS, amongst which, there have to be MTs minus-and plus-end-interacting proteins recruited for growing MTs tips (Sánchez-Huertas and Lüders, 2015; Paz and Lüders, 2018). Therefore, to give yolk asters a molecular identity as MTOCs, we analysed the expression of three different proteins (EB3, ⍰-tubulin and ß-tubulin).

*Eb3-mCherry* mRNA was injected at one-cell stage dclk2-GFP embryos to label MT plus-ends (Stepanova *et al*., 2003). Its recruitment into growing MT tips was used as an indicator of the polymerization process during plus-end growth occurring at the yolk MTOCs. Thus, e-YSN centrosomes were marked by EB3 comets as newly polymerized plus ends that emanate from them (Fei *et al*., 2019) and extended further down along growing MTs (Figures 4A and 4D). EB3 localized to e-YSN centrosomes and the mitotic spindle throughout mitosis, as shown in Video S7. Interestingly, EB3 comets also spread towards the vegetal pole throughout the yolk cell and far from YSN centrosomes (Figures 4B, 4C, 4E and 4F) which could indicate the non-centrosomal origin of some MTs.

**Figure 4.**
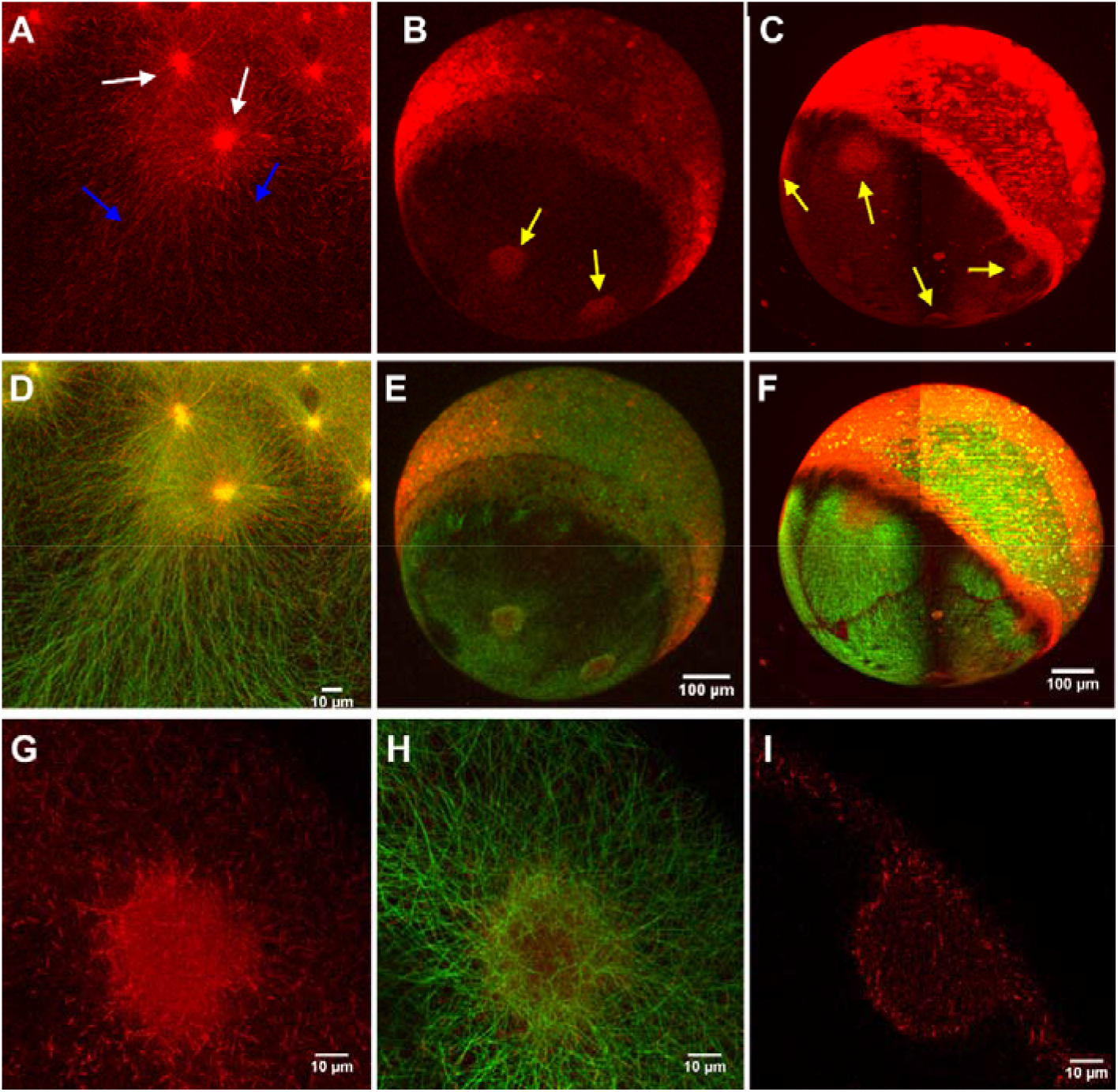
MT polymerization occurs at YSN centrosomes and YCL asters. *EB3-mCherry* injected dclk2-GFP embryos allows us to simultaneously visualize MTs (green) and MTs plus ends (red). EB3-mCherry signal was found: (A) in high concentration around e-YSN centrosomes (white arrows), as vegetalward oriented tracks in the YSL (blue arrows) and scattered puncta all along the entire yolk cell. See also Video S7. Whole imaging of the embryos using **(B)** LSCM and **(C)** LSFM also reveals an increase of signal at YCL asters at later stages (yellow arrows). (D-F) Merge of the EB3-mCherry and dclk2-GFP signals. **(G-H)** High resolution imaging and **(I)** cross section of a YCL aster shows the MT polymerizing activity at those MTOCs.

In the case of the YCL asters, a high density of EB3-mCherry signal was found at the core of every aster (Figures 4B, 4C, 4E and 4F), suggesting an increase of MT polymerization activity. Zoom into those asters (Figures 4G and 4H) provides a picture of the distribution of EB3 puncta along MTs, with higher signals at the YCL aster core. The cross-section of each of these centers reveals that EB3 distributes mainly in two opposed thin layers separated by a space with lower density of EB3 signal (Figure 4I), very similarly to the MT arrangement as shown in Figure 1I.

⍰-tubulin is a MT end protein identified as one of the key nucleator components of MTOCs (Wiese and Zheng, 2006). Although also found in nc-MTOCs, its function there as MTs nucleator, capper or stabilizer is not yet clear (Teixidó-Travesa, Roig and Lüders, 2012; Sanchez and Feldman, 2017). An immunostaining against ⍰-tubulin protein (Figures 5A-C) allows confirming that both blastoderm (Figure 5B) and e-YSN centrosomes (Figure 5C) contain ⍰-tubulin. At later stages, when YCL asters are formed, the injection of ⍰*tubulin-TdTomato* mRNA at one-cell stage dclk2-GFP embryos (Figures 5D-F) reveals high level of ⍰-tubulin protein at the core of every YCL aster, suggesting that yolk asters have a nc-MTOCs identity.

**Figure 5.**
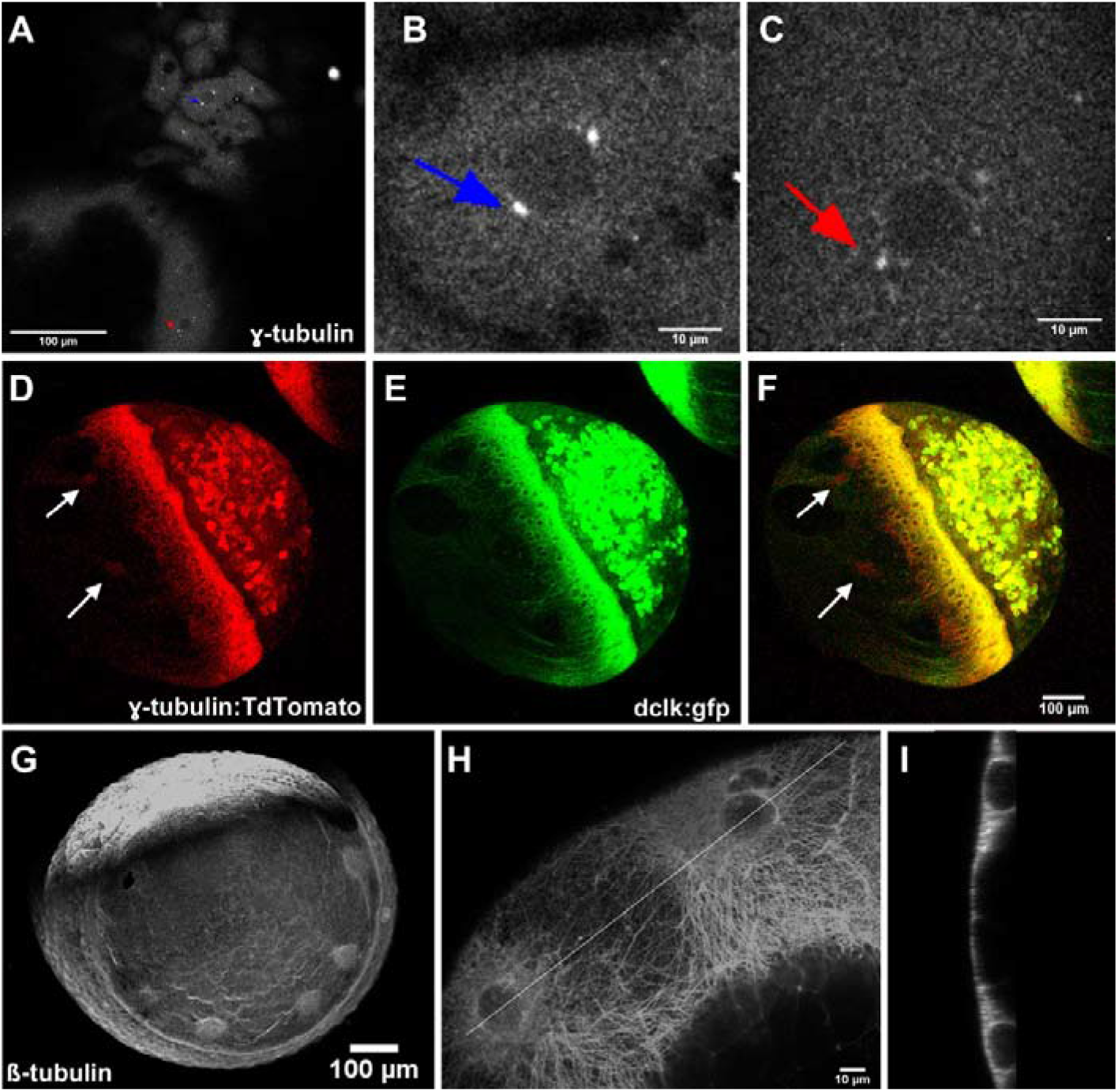
MT nucleation occurs at YSN centrosomes and YCL asters. ⍰ -tubulin expression can be observed through (A-C) immunostaining on fixed embryos and (D-F) ⍰-*tubulin-TdTomato* mRNA, coinjected with *dclk2-gfp* mRNA. Signal has been found in (B) blastoderm centrosomes (blue arrow), (C) e-YSN centrosomes (red arrow) and (D and F) YCL asters (white arrows). (G) ß-tubulin protein is detected in MTs and in high density in YCL asters (H). (I) Re-slice of two neighboring YCL asters shows that ß-tubulin fills each aster 3D volume. Images obtained through LSCM.

An immunostaining against ß-tubulin highlights MTs and MTOCs, revealing high concentration of ß-tubulin in the YCL asters, a sign of tubulin polymerization on those sites (Figures 5G-H). ß-tubulin fills in the YCL aster volume, in a 2-layered distribution (Figure 5I), similarly to dclk and EB3 proteins (Figures 1I and 4I, respectively).

In conclusion, the localization of EB3, γ-tubulin and ß-tubulin indicates that not only e-YSN centrosomes, but also YCL asters are sites of MT nucleation and growth.

### Centrin highlights centrosomal and noncentrosomal MTOCs in the yolk cell

To further characterize the molecular nature of the different yolk MTOCs we transiently expressed a centrin-GFP construct by injection of the corresponding mRNA into wild type embryos at one cell stage. We observe a diffuse signal in the nucleus and the cytoplasm of the blastoderm cells and the YSL, as well as brighter concentrations on both cells and e-YSN centrioles and other particles around the yolk membrane. This allowed to directly visualize, for the first time to our knowledge on zebrafish embryos, the centrosomic centrin that organizes the YSN mitotic spindles (from now on referred to as *Centrosomal Centrin*, CC) (Figures 6A and 6C). We found out that the same structures also contain ⍰-tubulin, highlighted with an antibody against the endogenous protein (Figures 5A-C, 6B and 6C). Interestingly, we discovered an independent second set of centrin within the YSL not associated to YSN (from now *on Anchoring Centrin*, AC), to anchor the AV parallel MT arrays (Figure 6D). Finally, a third set of centrin (from now on *Flowing Centrin*, FC), dispersed along the yolk cytoplasm, flows towards the animal pole along the MT network towards the YSL (Figure 6E). This retrograde flow takes place from blastula stages and continues even after YSN have become post-mitotic (Video S8). In few embryos, we also found a higher concentration of centrin signal all over the hemisphere with some distinguishable, highly dynamic, dense centers (Figure 6F). Remarkably, these centers resemble, in terms of size, morphology and dynamics (Video S9), the YCL asters observed in the Tg dclk2-GFP embryos. Although unusual, it is sometimes possible to find asters in wt embryos. This suggests that YCL asters may also contain an aggregation of centrin molecules. We tried, unsuccessfully, to develop a red version centrin tag in order to verify our observations on dclk2-GFP embryos. In any case, the identification of these four different subsets of centrin protein (CC, AC, FC and within the YCL asters) reveals the high complexity of the yolk MT organization.

**Figure 6.**
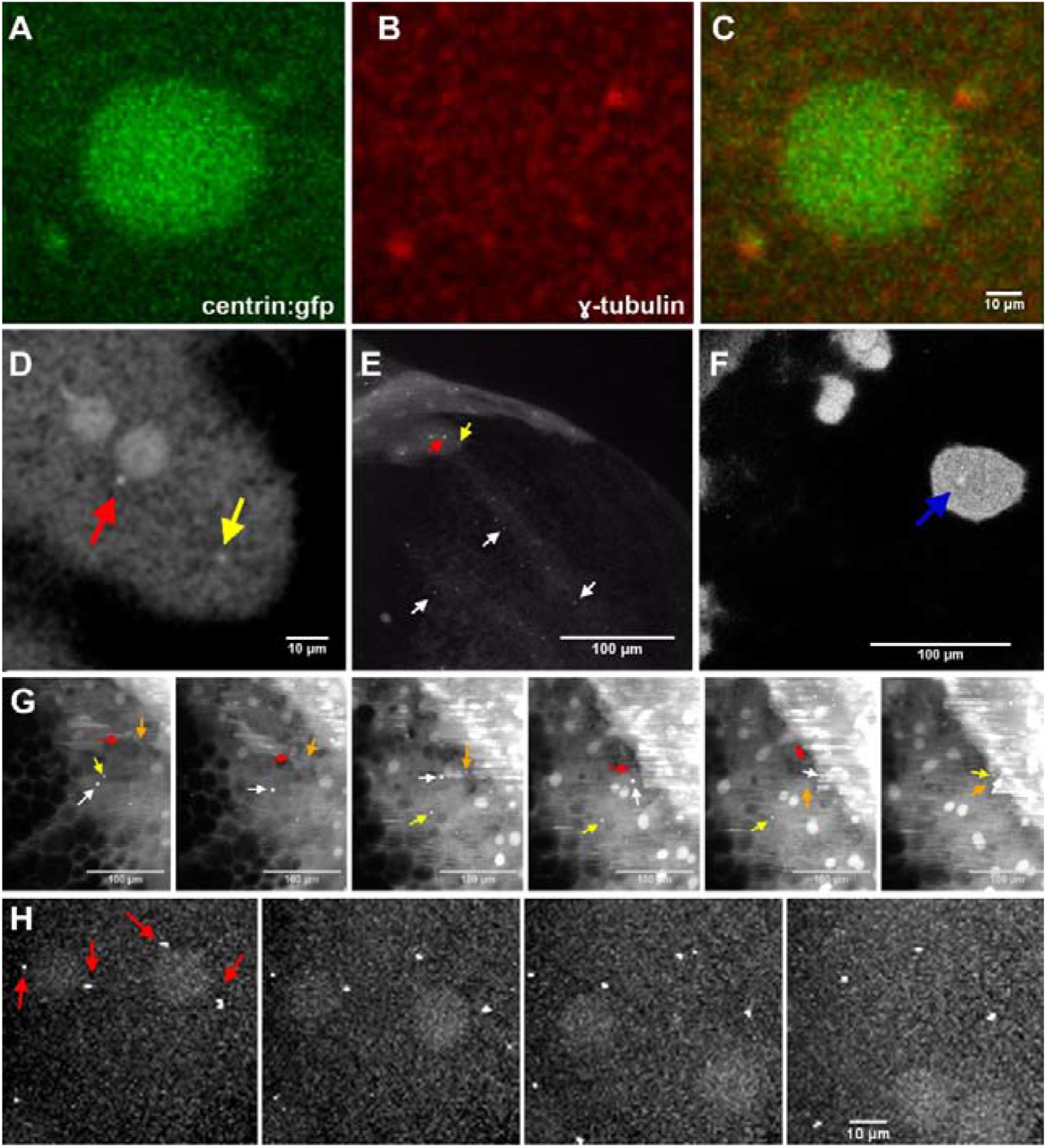
Centrin highlights centrosomal and non-centrosomal MTOCs in the yolk cell. **(A)** Centrin-GFP is expressed in e-YSN centrosomes and as a diffuse signal in e-YSN nuclei. (B) In comparison, ⍰-tubulin is only found at e-YSN centrosomes. **(C)** Merge of centrin and ⍰-tubulin signals. High centrin concentration can also be found at: **(D)** the YSL but not associated to YSN (*Anchoring Centrin*, AC), from where the AV MT arrays emerge (yellow arrows); (E) the yolk membrane, distributed as scattered puncta (white arrows), that flow animalwards, following MT arrays (*Flowing Centrin*, FC); and (F) the core of structures similar to the described YCL asters (blue arrow), visualized with LSCM. Also see Video S9, where centrin accumulation in the YCL resembles by shape, dimensions and dynamics, dclk2-GFP asters. **(G)** Here, zoom into the YSL region, where the interaction between FC (white arrow), AC (yellow arrows) and CC (red and orange arrows) suggests MTOCs functional reassignment by centrin re-localization during epiboly. Video S10 shows FC moving animalwards and its interaction with AC and CC, while epiboly progresses. Tiles in **(G)** are snapshots of a time lapse. **(H)** High resolution LSCM visualization of the YSN migration process initiation. The e-YSN are visible by a centrin diffuse signal and start migrating when centrosomes (red arrows) dettach. Tiles in (H) are snapshots of a time lapse.

The simultaneous visualization of all those different components through LSFM, allows us to describe a systemic centrin behaviour along the whole embryo, and specially the yolk cell, during all the epiboly process. As blastoderm and YSN divide, centrioles and nuclei are clearly visible allowing the tracking of mitotic processes. Interestingly, e-YSN centrioles remain after e-YSN exit the cell cycle. Afterwards, when e-YSN stop dividing, many bright FC puncta migrate from the vegetal pole towards the YSL following determined pathways, most likely streams of highly concentrated yolk MTs. These vesicles show a dynamic behaviour, with a tendency to aggregation. Surprisingly, we observed how those vesicles scan the YCL and interact with the e-YSN centrioles (Figures 6G and Video S10). We could clearly observe that at a specific moment, centrioles seem to jump out/detach of the YSL membrane. After the release of the centrioles, e-YSN start a fast migration towards the vegetal pole of the embryo (Figure 6H). Travelling FC puncta as well as CC migrate underneath the blastoderm as e-YSN migrate vegetalwards. As epiboly progresses and once e-YSN dissociate from their centrioles, the latter seem to be free to move within the YSL and/or actively pulled away by forces applied by the YSL cytoskeleton. In some cases, e-YSN centrioles seem to be dragged together with the traveling FC puncta and “disappear” beneath the blastoderm, within the internal-YSL (i-YSL), where we suspect they also undertake a role to organize the i-YSL MT network.

### The yolk MT network adopts different configurations in relation to the expressed levels of MT nucleating proteins (DCLK and DCX)

Next we sought to understand the mechanism responsible for the variability observed in dclk2-GFP embryos yolk MT organization. We combined data from our high-throughput LSFM microscope with brightfield inspection in order to screen tens of dclk2-GFP transgenic embryos. We confirmed that the number of YCL asters amongst eggs laid by different females differ in number, although it is on average repeated across different layings of given females. We have identified at least 8 females with offspring displaying this particular phenotype, and calculated the average number of YCL asters that the progeny of a specific female presents. This ranges between 0 and 22, depending on the female under analysis (Figure 7A). The presence of YCL asters in the dclk2-GFP transgenic line is transmitted across generations and therefore does not prevent development, i.e. eggs develop and become fertile adults.

**Figure 7.**
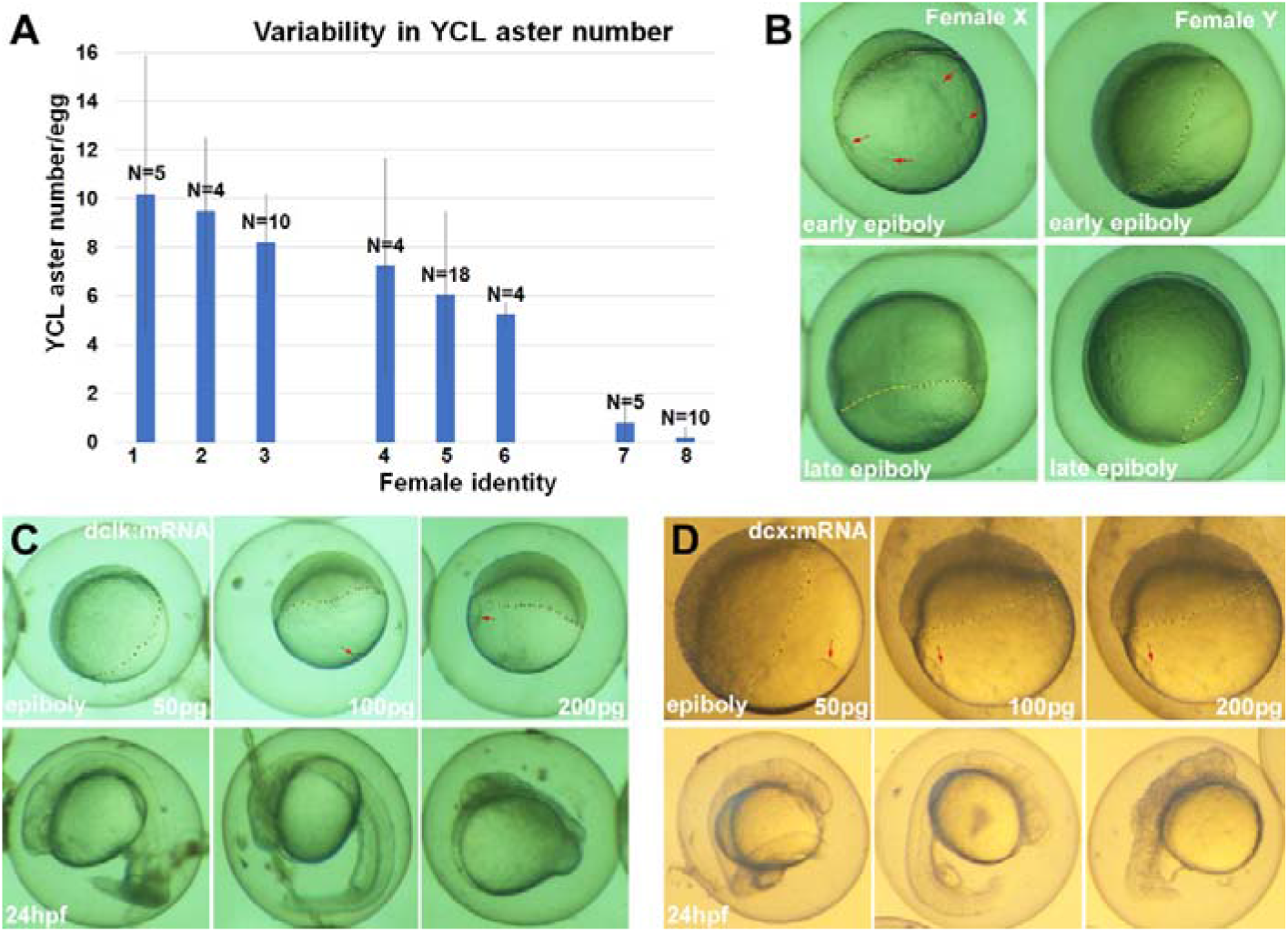
Natural and induced variability in YCL aster features and their impact on early development. **(A)** Comparison between the average number of YCL asters in eggs of selected Tg dclk2-GFP females: ≥8 YCL asters (females 1-3), between 3 and 7 YCL asters (females 4-6),≤2 YCL asters (females 7-8). N stands for the total number of eggs analyzed for the different females. **(B)** Brightfield imaging on representative sibling embryos from different Tg dclk2-GFP females: female X, medium number of asters, (10 eggs analyzed) and female Y, low number of asters, (10 eggs analyzed) at early (upper row) and late (bottom row) epiboly stages (epiboly extension: dashed yellow line. YCL asters: red arrows) **(C)** *dclk2-gfp* mRNA injections partially recapitulate the transgenic phenotype. Upper row: increasing mRNA doses proportionally induce a delay in epiboly (dashed red line) (Wt embryos, N=30; 50pg-injected embryos, N=20; 100pg-injected embryos, N=35; 200pg-injected embryos, N=20). Ectopic asters (red arrows) are formed from 100pg dose. Lower row: development follow up. (D) Equivalent experiment with *DCX-gfp* mRNA injections produces bigger asters (upper row, red arrows) and developmental defects (lower row) at lower doses compared to experiment in **(C).**

To understand if the different yolk MT phenotypes had an impact on epiboly progression, we monitored epiboly extension of the progeny of different transgenic fish. We found that epiboly is slower in dclk2-GFP eggs with more YCL asters. While wt embryos (N=10) and embryos (N= 10) with low number of YCL asters were at 50% epiboly (100%), their sibling embryos (N=12) with many YCL asters were still at sphere-doming stage (100%), and this delay persisted throughout epiboly (Figure 7 B).

In an aim to find the molecular mechanism behind the different observed yolk MT configurations, and suspecting a correlation with different dclk protein levels, we also performed *dclk-gfp* RNA injections in wt eggs at one cell stage, at different amounts (50 pg, 100 pg, and 200 pg). 100pg *dclk-gfp* RNA dose is able to induce the formation of YCL asters, that become bigger at 200pg dose. Already from 100pg dose we observe a delay in epiboly progression: while the wt embryos (N=30) and their 50pg-injected siblings (N=20) are at 60% epiboly (100%), the 100pg (N=35) are at 50% epiboly (40%) and 30% epiboly (60%), and 200pg-injected siblings (N=20) are at 30% epiboly (100%). Finally, 100pg injected embryos develop a shorter axis at 24hpf and 200pg dose produces body malformations and a complete lack of body axis extension in the most severe cases (Figure 7C).

Interestingly, asters can also be produced in the presence of low levels (50pg) of the paralog gene doublecourtin (DCX), by the injection of *DCX-gfp* RNA at one cell-stage. The overexpression produces big asters in the YCL that resemble those obtained in the *dlck-gfp* RNA injected embryos. They also display a dose-dependent delay in epiboly and developmental defects apparent at 24hpf (Figure 7D).

In conclusion, these experiments show that increasing the levels of these two closely related members of MT nucleating proteins family creates new scenarios. We observe that the yolk MT network, normally uniformly covering the yolk cell can re-organize into a multi-aster MT network. As long as the protein levels are kept in an appropriate range (stable transgenic line or low dose injection of corresponding construct) the phenotype is compatible with development, at this spatio-temporal window of observation. This represents an example of plasticity but robustness in development and indicates that this domain has the potential, both molecularly and biophysically, to change and adapt.

## DISCUSSION

A MT is a self-organization system in which tubulin monomers organize into dynamic filaments. A next level of complexity, i.e. the generation of a 3D pattern such as an aster, involves nucleation, filament dynamics and molecular motors (Nédélec, Surrey and Karsenti, 2003). *In vitro* reconstitution experiments show the formation of individual asters in polymer-stabilised microfluidic droplets, by the molecular motor-mediated contraction of a spherical network (Juniper *et al*., 2018). *In vivo*, a radial rearrangement of MTs in fish melanophore fragments is achieved through dynein-dependent MT nucleation (Vorobjev, Malikov and Rodionov, 2001). The aster-type pattern allows to explore the intracellular space and to define the position of the organelles through interaction with MT motor proteins. In this work, we discover MT asters in the zebrafish YCL and thus we update the current knowledge of zebrafish yolk MT organization. Despite the fact that it has been shown that yolk AV parallel MTs arrays are essential for the epiboly of the e-YSN (Solnica-Krezel and Driever, 1994; Fei *et al*., 2019), it turns out to be difficult to assess specifically the function of YCL asters. Further YCL asters interference experiments by locally applying nocodazole in the yolk or by uncaging a photoactivatable derivative of the MT depolymerising drug combretastatin (Wühr et al., 2010; Costache et al., 2017) could contribute to explain YCL asters role. Preliminary efforts trying to confine the drug effect in the YCL asters are being conducted. We hypothesise that they are used for mechanical and structural support of the huge zebrafish yolk cell during epiboly, and that they form via a combination of MT nucleation and molecular motors confined in the spherical embryo. The profound plasma membrane deformations identified with bright field microscopy (Figure 1G) indicate potential MTs anchoring sites, providing enough mechanical force to induce those deformations. In these sites, YCL asters would be serving for mechanical support. Indeed, it is known that cytoskeletal elements can pull membranes by polymerization or with the help of motor proteins (Solinet *et al*., 2013; Jarsch, Daste and Gallop, 2016).

Comparing to previous literature, the new array of yolk MTs that we describe in this work is different from the AV oriented YCL longitudinal MTs that e-YSN use to migrate vegetalwards (Solnica-Krezel and Driever, 1994; Fei *et al*., 2019). In many embryos there is a clear MT void space between the AV MT arrays emerging from the blastoderm and the domains formed by the YCL asters. In fact, we found that e-YSN migration along the YCL longitudinal MTs occurs concomitantly with the presence of these vegetal YCL asters (Figures 3K and 3L and Video S5). In any case, until the MT network is re stablished, during YCL aster reabsorption, e-YSN migration seems to be paused, confirming the need of the AV MT network in order to move forward.

Despite the work done with the reconstitution of multiple large asters in a cell-free system (Mitchison *et al*., 2012; Ishihara *et al*., 2014; Ishihara, Korolev and Mitchison, 2016), to our knowledge there is no report of live non-artificial models forming big multiple MTOCs in an individual cell, with the exception of the transient formation of several small MTOCs in the immature mouse oocyte (Schuh and Ellenberg, 2007; Li and Albertini, 2013). Our data supports that multiple YCL asters form naturally in many of the analyzed eggs, and therefore the zebrafish yolk cell arises as a promising natural model where to analyze aster-aster interaction zones (Figure 3I). In our model, these regions are very similar to the boundary zones between non-sister asters following polyspermic fertilization in amphibian embryos. Most probably, YCL aster growth is limited by the presence of the neighboring aster, where anti-parallel MTs overlap (Mitchison *et al*., 2012). In fact, previous extensive studies have reported that Aurora B kinase is key for the formation of asters boundaries in *Xenopus* extracts and between asters in the midplane of cleaving Xenopus zygotes (Field, Pelletier and Mitchison, 2019). How similar our boundary zones are to other studied boundaries, both biophysically and molecularly, remains to be examined.

Moreover, we observe an almost perfect synchrony between the multiple YCL asters, equidistant to the YSL, across the different steps leading to their formation and re-absorption, pointing to 1) a global coordination mechanism and to 2) a tight mechanical dependence on the epiboly process. The radial distribution in concentric rings around the AV axis without any apparent dorso-ventral bias, also supports this last idea, since epiboly is essentially a radially symmetrical process in this direction. We found that YCL asters are not randomly located in the embryonic sphere. On the contrary, they tend to group in the same latitude (or various latitudes when more than one ring is formed) and to keep a quasi-constant distance between them (Figure 2). We suggest that this distribution is related to geometrical or mechanical constrains. The YCL asters resemble polygon-like structures that cover the exposed yolk surface (see figure 2A) (it remains to be tested if the non-exposed, i.e underneath the blastoderm, is also covered by YCL asters). Thus, remarkably, the yolk is revealed as a biological model for the uniform distribution of many points in a (hemi)sphere. Accessing the totality of the yolk sphere maybe by mechanical removal of the blastoderm would allow to test if the yolk MT network is a solution in nature for the classical best packing problem in spheres and minimal energy point configurations (Saff and Kuijlaars, 1997; Brauchart and Grabner, 2015).

In this work we prove that e-YSN spindles are organized through MTOCs, by the localization of EB3, ⍰-tubulin and centrin proteins. Our observations agree on the experiments conducted by (Fei *et al*., 2019) where e-YSN centrosomes are highlighted with an EB3-tagged construct. We also show that the presence of EB3 and ⍰-tubulin in YCL asters unequivocally indicates that they are sites of MT nucleation (Figures 4 and 5). Finally, this is supported by the localization of high levels of β-tubulin, used for MTs assembly, in the asters (Figures 5G-I).

In fact, LSCM and LSFM have been instrumental for the localization and dynamics of the different MTOCs components and to understand that several MTOCs need to be in place to organize a network of such dimensions, that dramatically changes over the early stages of zebrafish development. In particular, we prove that transient expression of centrin-GFP protein (CC, AC and FC -see section 4 of Results) successfully highlights centrosomal and noncentrosomal MTOCs in the yolk cell, in agreement with (Paoletti *et al*., 1996) that stated that more than 90% of centrin is not associated with the centrosome fraction. The YCL FC moves animalwards, towards the YSL, and seems to follow the AV MT parallel arrays tracks (Video S8). Travelling FC puncta briefly interact with CC centrioles (Video S10) and migrate underneath the blastoderm as e-YSN migrate vegetalwards. We suggest that this FC-CC centrin interaction is probably part of the process that actually triggers the e-YSN migration. Interestingly, e-YSN centrioles disassociate from these nuclei, most probably through a detachment mechanism from the nuclear envelope, and become “free” to drift away from the nuclei (Figure 6) (Archambault and Pinson, 2010). From the stage e-YSN exit the cell cycle, we assume that their centrosomes lose their activity as MTOCs, and new cellular sites are specified and activated to acquire this function within the huge yolk cell. In differentiated cells, the loss of centrosomal MTOC activity and the formation of non-centrosomal sites can happen through different mechanisms (Keating *et al*., 1997; Bartolini and Gundersen, 2006; Brodu *et al*., 2010; Muroyama and Lechler, 2017; Sanchez and Feldman, 2017) although most studies point to protein localization and pericentriolar material delocalization as the most common ways of regulation. However, in our system we hypothesize that the mechanism is slightly different, because the formation of the novel more vegetal yolk MTOCs does not imply the disappearance of the e-YSN CC (Figure 6). Concomitantly, yolk MT network organization seems to be governed by several MTOCs elsewhere, rather than only the centrosomal-MTOCs (e-YSN CC). This would imply the recruitment of centrin and/or other key molecules to those new sites probably through a MT and motor proteins-dependent mechanism (Dammermann and Merdes, 2002). Centrin behaviour in our model supports this idea: first, we observe that e-YSN CC seems to be recycled, because e-YSN centrioles do not disappear. Second, at middle to late epiboly stages, we observe centrin expression in structures located more vegetally, resembling YCL MTOCs both by location and behavior. Thus, centrin is at the core of both centrosomal and non-centrosomal yolk MTOCs that co-exist in the embryo. The observed centrin temporal and spatial dynamics inform on the biology of centrosomes and points to a complex whole-embryo large-scale regulation that deserves future study.

It has been proposed that asters can form in different ways either in natural or artificial conditions (Nédélec, Surrey and Karsenti, 2003). We have specifically investigated the dclk2-GFP transgenic line and found that it represents a scenario in which the phenotypic variability (absence/presence and different number of asters) accessed through LSFM could reflect the different levels of dclk protein expressed in the progeny of different tg dclk2-GFP individuals (Figures 7A and 7B). Importantly, this provides evidence that the yolk MT organization doesn’t follow a rigid scheme. We propose that the variability observed is a possible outcome of the use of the Tol2 transgenesis method (Kawakami, 2007), which could have produced different integration sites of the Tol2 element in the respective genomes. This suggests that our model of study is the result of an overexpression phenotype (Prelich, 2012) compatible with normal development as long as the gene is expressed at an appropriate range level. Thus, *dclk2* or *DCX* mRNA injections are not neutral at higher doses (Figures 7C and 7D), while the transgenic line is stable and all analysed phenotypes are compatible with development, probably because expression levels are kept within an appropriate range. We speculate that increasing the levels of DCLK or DCX causes the formation of ectopic nucleation sites, in our case YCL asters, similar to ⍰-tubulin or RanBPM overexpressions (Shu and Joshi, 1995; Nakamura *et al*., 1998). However, the transient expression induces the uncontrolled formation of YCL asters that are bigger and misslocalized when compared to the asters formed in the stable transgenic line, giving rise to developmental defects.

We rarely found YCL asters in wt “naked” eggs (non-injected, non-transgenic), which could be explained in two ways. On one hand, despite YCL asters might exist, the expression levels of the molecules recruited to those sites are too low to be detected with our tools and to generate big asters capable to induce the profound yolk membrane deformations observed with bright-field imaging (Figure 1G). On the other hand, it is possible that wt embryos organize the yolk MT network in an alternative way and YCL asters are an example of the creation of a functionality that is normally hidden in the system, that is triggered by the overexpression of MT nucleating proteins in a particular time and territory.

With this work we have specifically investigated the organization of the yolk MT network during zebrafish epiboly. The transgene-mediated phenotypic variability that we have found points to the yolk domain as a plastic but robust territory, able to present diverse configurations. This variability however encloses universal mechanisms for MT organization, especially in large cells, like the zebrafish yolk cell, where different MTOCs are in place to ensure development. Finally, our results underscore the importance of the observation of the embryo as a whole, allowing to connect events in time and in space.

## METHODS

### ZEBRAFISH STRAINS

AB and Tg:(XlEef1a1:dclk2a-GFP) strains (from Marina Mione, CIBIO, University of Trento, Italy) (Tran, L.D.,et al., (2012) were used. Animals were housed under standard conditions.

Zebrafish embryos were kept in E3 medium (Nüsslein-Volhard and Dahm, 2002) and staged as previously described (Kimmel and Law, 1985). Embryonic manipulations were done in E3 medium. The embryos analyzed in our study are always the result of outcrosses between Tg:(XlEef1a1:dclk2a-GFP) females with AB WT males. Up to 15 Tg:(XlEef1a1:dclk2a-GFP) females of successive generations were used in this work.

### DNA CONSTRUCTS AND mRNA INJECTIONS

The following expression constructs were used: EB3-mCherry (Stepanova *et al*., 2003), centrin-GFP, ⍰-tubulin-tdTomato, (kindly provided by Virginie Lecaudey, Goethe-Universität, Frankfurt), dclk2-GFP (kindly provided by Marina Mione, CIBIO, University of Trento) and DCX-GFP (kindly provided by Esteban Hoijman, CRG, Barcelona). mRNAs were synthesized using SP6 mMessage machine kit (Ambion^®^, Life Technologies, Germany), after NotI linearization. Zebrafish embryos were injected using glass capillary needles (Harvard apparatus 30-0020 GC100F-15) which were pulled with a needle puller (Sutter P-97), and attached to a microinjector system (World Precise Instrument PB820). Unless otherwise indicated, 100 pg of EB3-mcherry mRNA, centrin-gfp mRNA, dclk-gfp mRNA or ⍰-tubulin-tdTomato mRNA were injected into 1-cell stage embryos.

### WHOLE-MOUNT IMMUNOHISTOCHEMISTRY

Mouse anti-β-tubulin antibody (E7, Developmental Studies Hybridoma Bank, DSHB) was used at 1:200 and mouse anti-γ-tubulin antibody (T5326 Sigma-Aldrich) was used at 8ug/ml. The secondary antibody was in-house conjugated (Bálint *et al*., 2013) to the Abberior STAR 635P fluorophore (Sigma) and used at 8ug/ml.

γ-tubulin staining was performed as previously described (Li-Villarreal *et al*., 2015) Briefly, embryos were fixed in 4%PFA overnight at 4ºC. Fixed embryos were washed several times in PBS and dechorionated. Permeabilization was done in 0.3% Triton X-100 (in PBS) for 1 h, exchanging buffer every 15’. Blocking was performed in blocking solution (1% BSA in 0.3% Triton X-100 in PBS) for 2 hours, at RT. Embryos were incubated with γ-tubulin primary antibody in blocking solution overnight at 4ºC. After washing the embryos in 0.3% Triton X100 in PBS overday, they were incubated in the secondary antibody in blocking solution overnight at 4ºC. After 3-4 washes in PBS, the embryos were ready to be mounted and imaged.

ß-tubulin antibody staining was performed as previously described (Topczewski and Solnica-Krezel, 1999) with some modifications. Briefly, embryos were dechorionated and fixed in MT assembly buffer (80 mM KPIPES (pH 6.5), 5 mM EGTA, 1 mM MgCl2, 3.7%formaldehyde, 0.25% glutaraldehyde, 0.5 uM taxol, and 0.2% TritonX-100) for 6 hours at RT. Fixed embryos were dehydrated and kept in methanol at −20°C overnight, or for several days. After, they were washed several times in PBS containing 0.1% NP40, for re-hydration. Re-hydrated embryos were then incubated in 100mM NaBH4 in PBS for 6-16 hours at RT, and washed extensively in tris buffered saline (TBS). Blocking was performed in blocking solution (2% BSA in TBS) for 30’ at RT. Embryos were incubated with β-tubulin primary antibody in blocking solution overnight at 4ºC. After washing them 4-5 times in TBS, they were incubated in the secondary antibody in blocking solution for 2-3 hours at RT. After 3-4 washes in TBS they were ready to be mounted and imaged.

### IMAGING SET UPS AND SAMPLE PREPARATION FOR LIVE AND FIXED IMAGING

LSCM was performed on a commercial Leica SP8 equipped with a supercontinuum white-light laser (KTT) and Hybrid detectors, and with HC PL APO CS2 10x/0.40 DRY and HC PL APO CS2 63x/1.40 OIL objectives. For live imaging, dechorionated embryos were mounted in 0.5% low melting point (LMP) agarose (ref) in E3 medium, on glass bottom dishes (MatTek). Fixed samples were mounted in 1% LMP agarose in PBS, on glass bottom dish (MatTek).

LSFM imaging was performed using a custom-made light-sheet set up, an improved version of our previous design (ref), called Flexi-SPIM (ref). For illumination we used 488 and 561 nm lasers (Cobolt, MDL488 and MLD561) and two air objectives (Nikon 4x, NA0.1). For fluorescence detection we used water dipping objectives (Nikon 10x, NA0.3 and 20x, NA 0.5), filters and a sCMOS camera (Hamamatsu OrcaFlash4 v2). Thanks to our configuration, multiple views of the specimen can be visualized, providing in toto embryo representations. We realized, along the experiments, that LSFM imaging of the yolk leads image degradation along the illumination axis. This is due to light refraction on the lipid filled spherical yolk cell, that acts as a lens. Sequential side illumination, although requiring a simple fusion process, increases image quality. However, this leads to a blind central region on the data sets. This effect is neglectable in the blastoderm. Embryos were simply mounted, without removing its chorion, within either a 1.5% LMP agarose cylinder or a 1mm inner diameter FEP tube filled with E3 medium.

Bright field images were acquired with a scope or the light-sheet set up and a source of white-light, bulb or led, respectively.

### IMAGE ANALYSIS

To evaluate YCL asters sizes, the 3D z-stacks were analysed. YCL MT asters’ depth and diameter were measured with FIJI in their max value.

For the quantification of the YCL asters number, we manually counted the number of YCL asters on the Z maximum projection of LSFM and LSCM images. Only the eggs offering a vegetal view (the totality of the YCL asters could be accessed) and eggs offering a lateral view, in which radial symmetry could be assumed, were considered. The females were grouped in 3 classes: females producing eggs with many asters (≥8), females producing eggs with an intermediate number of asters (between 3 and 7) and females producing eggs with a very low number of asters (≤2). The analysis includes 8 females in total.

To estimate the local orientation of the MT bundles in YCL asters (N=9) at different distances from the YSL, we used two Fiji plug-ins: OrientationJ (Püspöki et al., 2016) and Directionality (created by Jean-Yves Tinevez, Institute Pasteur). For OrientationJ we used in particular the functionality OrientationJ Analysis to render a visual representation of the orientation of the MTs in the YCL asters, and the functionality OrientationJ Vector Field to create a vector field map of the selected images. To generate the histograms that show the amount of MT bundles in particular directions the local gradient orientation method of Directionality plugin was used. Background was subtracted and a smooth filter was applied (FIJI) in the original images before the orientation analysis were performed.

To determine YCL MT aster distribution over the yolk, 3D stacks acquired either by LSFM or LSCM were analysed. Images relative to a single time point were used, correspondent to the embryo at 50-65% epiboly, i.e. when the YCL asters are clearly visible. A FIJI home-made macro permits to select and export the 3D coordinates of the centre of the embryo, the vegetal pole, and the asters. Elaborating these 3D coordinates through a MATLAB script, the AV axis orientation and the radius of the modelled yolk sphere are obtained. Based on this, a rigid body transformation of all the points is applied, so that the centre of the embryo is coincident with the coordinate origin, and the AV axis aligned along the “z” axis. Describing the YCL MT aster coordinates by mean of spherical coordinates, the latitude (polar angle relative to the AV axis) and the longitude (azimuthal angle) of each of the MT asters are computed. For each embryo, MT asters having a similar latitude (i.e. a maximum of 20 degrees of difference) were considered to belong to the same ring, concentric to AV axis. For each ring, the differences between longitudes of neighbour MT asters were calculated to examine eventual angular equidistance between them. Equidistance is defined as the angular distances’ differences is belonging to a 10 degrees range. A total of 11 embryos with at least 4 YCL MT asters were analysed. For angular equidistance analysis, only rings with more than 2 asters were considered. In cases where equidistance is not found, we suspect an aster in between could not be detected from the image 3D field of view.

## ACKNOWLEDGMENTS

This project and MB have received funding from the European Union’s Horizon 2020 research and innovation programme under the Marie Sklodowska-Curie grant agreement No 721537 “ImageInLife”. EG received funding from MINECO/FEDER Ramon y Cajal program (RYC-2015-1793). Authors also acknowledge financial support from the Spanish Ministry of Economy and Competitiveness through the “Severo Ochoa” program for Centres of Excellence in R&D (SEV-2015-0522), from Fundació Privada Cellex, from Fundació Mir-Puig, and from Generalitat de Catalunya through the CERCA program”. We thank Virginie Lecaudey, IZNF Germany, and Darren Gilmour, IMLS Switzerland, for the EB3-mCherry, centrin-GFP and γ-tubulin-TdTomato DNA constructs, Marina Mione, CIBIO, University of Trento, for the dclk2-GFP DNA construct and Esteban Hoijman, CRG Spain, for the DCX-GFP DNA construct. We are grateful to Timothy Mitchison for providing the combretastatin drug for preliminary assays. We thank Verena Ruprecht, CRG Spain, Jordi Andilla, ICFO, Spain, and Pilar Pujol, ICFO, Spain, for helpful discussions. We thank Senda Jiménez, CRG, Spain, all SLN-ICFO members and ICFO Biology lab members for their support.

## AUTHOR CONTRIBUTIONS

MM designed the experiments. MM, MB and EG conducted the experiments and analysis. MM and EG wrote the manuscript. MB and PL read and edited the manuscript. MM, and PL conceptualized the study. PL supervised the work.

## SUPPLEMENTARY INFORMATION

Document S1. Figures S1–S2

**Supplementary Figure 1:**
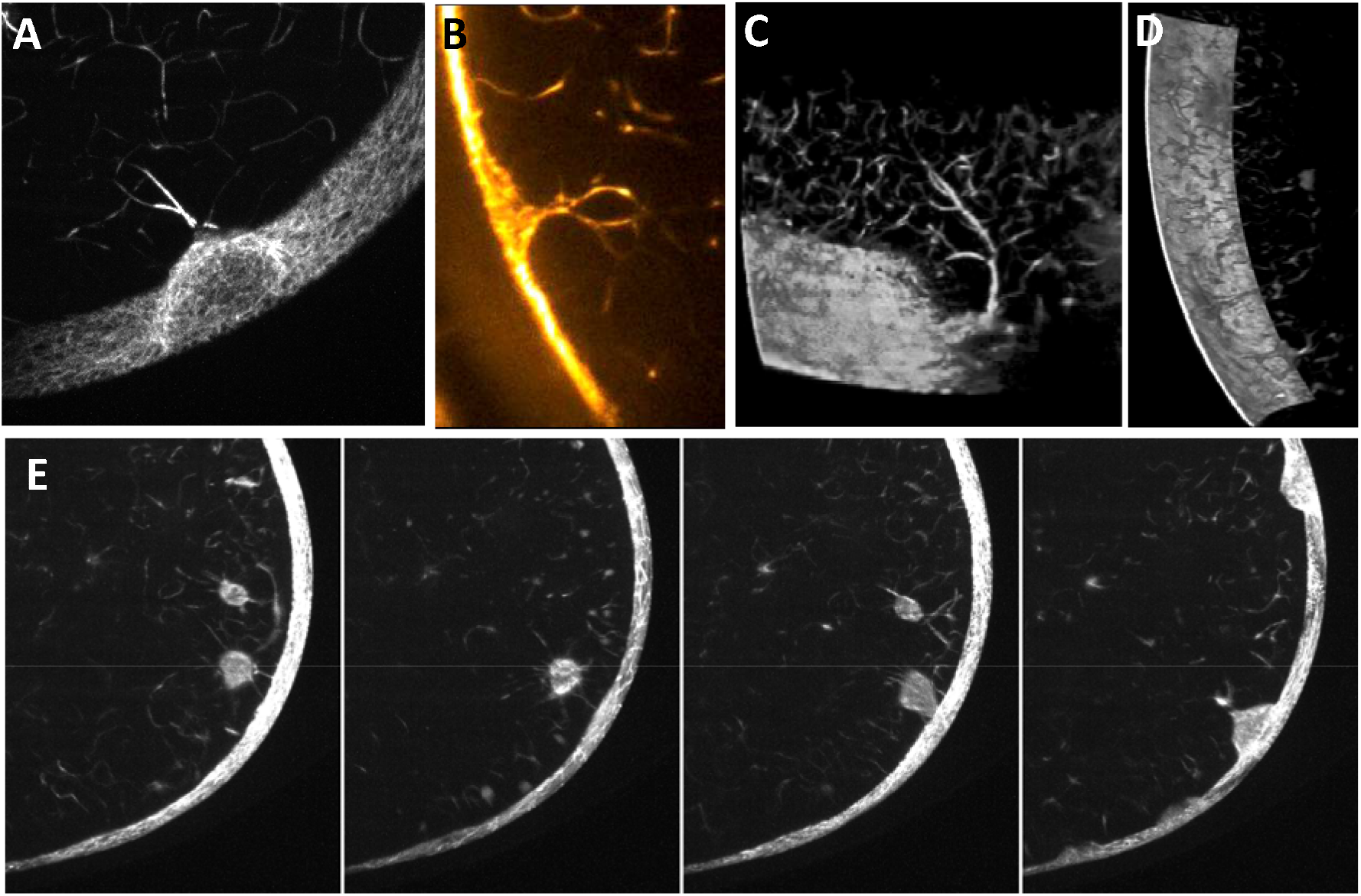
Light sheet fluorescence microscopy allows visualizing the inner yolk microtubule network and its interaction with YCL asters. **(A-D)** In some cases it is possible to observe that YCL asters are able to recruit the inner microtubule network forming dense MT bundles. (E) Observing over time a cross section of the embryo it is possible to observe how YCL asters recruit inner yolk MT aggregates. In any case, it is worthy to point out, that those examples do not fully represent the observed dynamics. Normally YCL asters form independently of its interaction with inner microtubules bundles or aggregates.

**Supplementary Figure 2:**
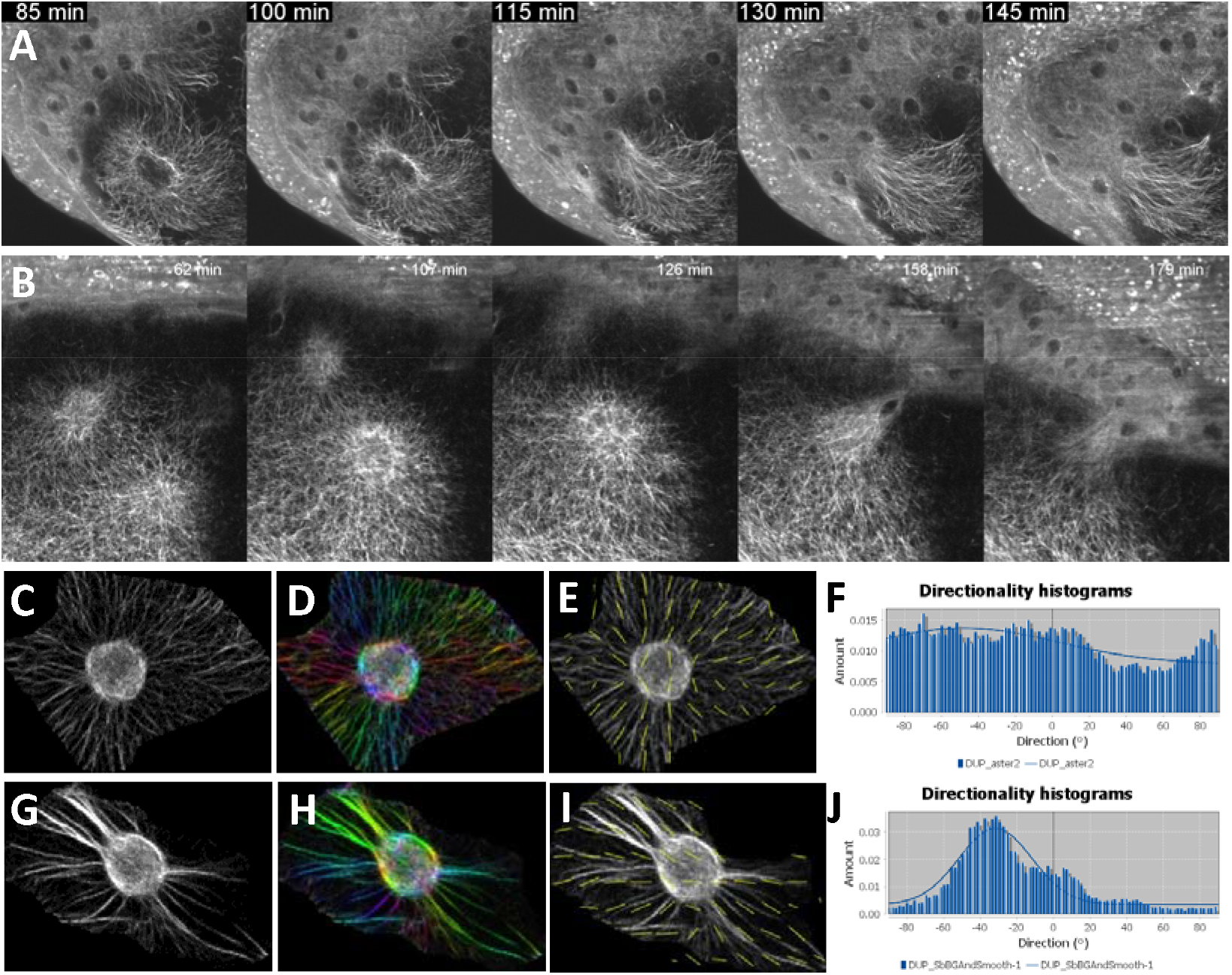
Orientation of the MT bundles within representative YCL asters regarding distance to blastoderm margin. **(A)** and **(B)** show two examples of YCL aster reabsorption as the blastoderm margin approaches. To estimate the local orientation of the MT bundles we used two ImageJ plug-in: OrientationJ and Directionality. **(C-F):** YCL asters not adjacent to blastoderm margin show an isotropic distribution of MT bundles. **(C)** Original image of YCL aster. **(D)** OrientationJ Analysis module performs a visual representation with a color map of the distribution of the MT bundles. (E) Vector field overlaid on the original image, performed by OrientationJ Vector Field module. **(F)** Histogram computed with Directionality plugin, indicating the amount of MT bundles in a given direction. This flat histogram indicates a very isotropic MT content in this type of YCL asters. (G-J) The same analysis was performed when the blastoderm margin approaches the YCL asters. (H) The histogram shows a distinguishable peak, indicating a preferred orientation of the MT bundles in the AV direction.

## VIDEO LEGENDS

Video S1. Video shows the three-dimensional render of half embryo with multiple YCL asters, Related to Figure 1.

Video S2. Video shows the MTs dynamics of the whole dclk2-GFP embryo during epiboly using Multi-view Light-sheet Fluorescence Microscopy, Related to Figure 2.

Video S3. Video shows an example of the formation of a vortex of MTs in the yolk of a dclk2-GFP embryo with a single YCL aster in the vegetal pole using Scanning Laser Confocal Microscopy, Related to Figure 2.

Video S4. Video shows the formation of a YCL aster in a dclk2-GFP embryo using High Resolution Scanning Laser Confocal Microscopy, Related to Figure 2.

Video S5. Video shows the reabsorption of YCL asters in a dclk2-GFP embryo using High Resolution Scanning Laser Confocal Microscopy, Related to Figure 2.

Video S6. Video shows the reabsorption of YCL asters in a dclk2-GFP embryo using Multi-view Light-sheet Fluorescence Microscopy, Related to Figure 2.

Video S7. Video shows EB3 (red) and MTs (green) dynamics during e-YSN division in a dclk2-GFP embryo using Scanning Laser Confocal Microscopy, Related to Figure 4.

Video S8. Video shows two examples of the animalwards flow and aggregation of centrin puncta in *centrin-gfp* mRNA injected wild type embryos using Multi-view Light-sheet Fluorescence Microscopy, Related to Figure 6.

Video S9. Video shows transmitted infrared (left) and GFP Fluorescence (right) time-lapse images of the YCL asters dynamics containing bright centrin puncta in a *centrin-gfp* mRNA injected wild type embryo using Scanning Laser Confocal Microscopy, Related to Figure 6.

Video S10. Video shows the animalwards flow and aggregation of centin puncta from the YCL and its interaction with e-YSN centrosomes, triggering the e-YSN vegetalwards migration, in *centrin-gfp* mRNA injected wild type embryos using Multi-view Light-sheet Fluorescence Microscopy, Related to Figure 6.

